# Seasonality and interspecific temporal partitioning in a semiarid grassland bat assemblage

**DOI:** 10.1101/2021.12.04.471220

**Authors:** Ram Mohan, Vaibhav Chhaya, Anand Krishnan

## Abstract

Arid and semiarid environments of the world are prone to dramatic seasonal changes that affect the availability of scarce, patchily distributed resources such as water. In response to these changes, animals migrate or partition resources to minimize competition, resulting in temporal patterns within assemblages across multiple scales. Here, we demonstrate that the winter dry season bat assemblage in a semiarid grassland of northwest India exhibits both seasonal changes in composition and temporal avoidance between coexisting species at water bodies. Using a passive acoustic monitoring framework to quantify activity patterns at different points in the season, we show that two species (*Rhinolophus lepidus* and *Tadarida aegyptiaca*) exhibit seasonal differences in activity, being more frequently detected in the early and late parts of the dry season respectively. Two other species (*Pipistrellus tenuis* and *Scotophilus heathii*) do not exhibit seasonal changes in activity, but structure their diel activity patterns to minimize temporal overlap (and thus competition) at water bodies. These data, some of the first on bats from this region, demonstrate the complex temporal patterns structuring bat assemblages in arid and semiarid biomes. Our results hold promise for monitoring efforts, as a baseline to ascertain how climate change may influence the behavior and ecology of desert and grassland organisms.

## INTRODUCTION

Forty-one percent of the earth’s surface area is occupied by arid and semi-arid habitats (Safriel and Adeel, 2008), characterized by extreme environmental conditions like high summer temperatures and low rainfall (Wickens, 1998). These environmental constraints drive the spatial and temporal distributions of animal taxa occupying these unique ecosystems. Wildebeest, for instance, migrate long distances in search of suitable grazing patches, and Arctic ground squirrels hibernate during the winter (Barnes, 1989; Bond et al., 2017). Extreme weather conditions may alter the distributions of important resources, particularly water. These seasonal changes influence the distributions of fauna occupying arid and semiarid environments, which depend on water resources (Smit and Grant, 2009; Van Heezik et al., 2003; Yosef and Zduniak, 2011). Monitoring the few water bodies that exist in these environments provides both an inventory of their fauna, as well as an indicator of spatial and temporal change. The scarcity of resources such as water may additionally lead to competition, particularly between closely related species (D’Onofrio et al., 2015). This, in turn, is predicted to increase avoidance between competing species, which manifests as spatial and temporal heterogeneity in the fauna detected at monitoring sites (Valeix et al., 2007). Monitoring seasonal patterns and movements of poorly-studied taxa provides critical information on their response to climate change, as well as inputs for conservation.

Among the many taxa inhabiting arid and semiarid ecosystems, volant animals such as bats present a unique opportunity to examine seasonal movement and activity patterns. Bats represent a high proportion of the mammalian diversity in arid and semiarid ecosystems (Carpenter, 1969), and are physiologically adapted to withstand climatic extremes (Marom et al., 2006; O’Farrell and Bradley, 1970). Multiple studies have added to our understanding of the seasonality of bat assemblages in temperate habitats. For example, migration and hibernation alter bat assemblage composition during cold temperate winters (Brooks, 2011; Nocera et al., 2019). Tropical regions with high summer temperatures experience changes in bat activity linked to the availability of perennial sources of water (Hagen and Sabo, 2012). These patterns demonstrate the sensitivity of bats to changes in local environmental conditions, thus rendering them excellent bioindicators to monitor change in arid environments. By monitoring and quantifying seasonality in bat assemblages, we may also understand how these patterns are affected by longer-term environmental changes.

As mentioned above, the scarcity of water in arid environments may also lead to competition at the few remaining sources, which in turn adds another dimension to the temporal structure of the assemblage. In an arid environment, where multiple bat species concentrate at a spatially limited resource like water, complex community-level interactions may take place (Adams and Hayes, 2021). Resource partitioning in sympatric bats can be observed along spatial, temporal and dietary axes (Beilke et al., 2021; Razgour et al., 2011). For instance, bats foraging over small water bodies, particularly species with similar ecological traits, may arrive at different times to minimize competitive overlap (Adams and Thibault, 2006; Cockrum and Cross, 1965; Lambert et al., 2018; O’Farrell and Bradley, 1970). This in turn has consequences for diel activity patterns and foraging activity within an assemblage. Monitoring bats at scarce water resources thus has the potential to inform our understanding of competition and community structure at multiple temporal scales, both diel and seasonal. In many of the world’s dryland ecosystems, particularly in the tropics, there is very little data on seasonality and structure of bat assemblages (Lisón et al., 2019). Specifically, do bat assemblages exhibit seasonal change as the dry season progresses? Does the availability of scarce resources such as water, and interspecific competition for these resources, drive temporal structure in bat assemblages? Addressing these questions helps us obtain a quantitative metric to measure how bat assemblages respond to climate change and its effects on the availability of water.

Here, we study seasonality and temporal partitioning in a bat assemblage occupying a semiarid grassland in Northwest India, at the edge of the Thar Desert. There have been very few studies of bat assemblages or communities in the Indian Subcontinent (Chakravarty et al., 2021; Ongole et al., 2018; Wordley et al., 2018), in spite of its high bat diversity, and virtually none on seasonality and temporal activity patterns around water bodies. Passive acoustic monitoring presents a powerful tool to study these questions (Lisón et al., 2019), especially given recent call libraries of bats from the region’s vicinity (Shah and Srinivasulu, 2020). We hypothesized that a) winter dry-season bat assemblages at water bodies should exhibit seasonal changes in structure, and b) that competing species should exhibit temporal avoidance to minimize overlap at scarce water resources. Our data, some of the first on the temporal structure of bat assemblages from the region, presents insights into interspecific interactions and seasonality in bats occupying dryland ecosystems, as well as a paradigm to study long-term responses of these species to climate change.

## MATERIALS AND METHODS

### Study area and sampling sites

We conducted this study in the Tal Chhapar Blackbuck Sanctuary, Churu District, Rajasthan (27.798490 N, 74.434921 E) (Figure 1). This small (approximately 7 square kilometres) sanctuary is a semiarid savanna grassland of the *Dichanthium-Cenchrus-Lasiurus* type (Dabadghao and Shankarnarayan, 1973) at the edge of the Thar desert (Kaur et al., 2020) with scattered bushes of *Acacia, Capparis, Calotropis* and *Zizyphus*, and water bodies which are maintained by the forest department. Our recording sites fell broadly into two categories - ***water bodies*** (three sites) and ***grassland*** sites (four sites).

**Figure 1:**
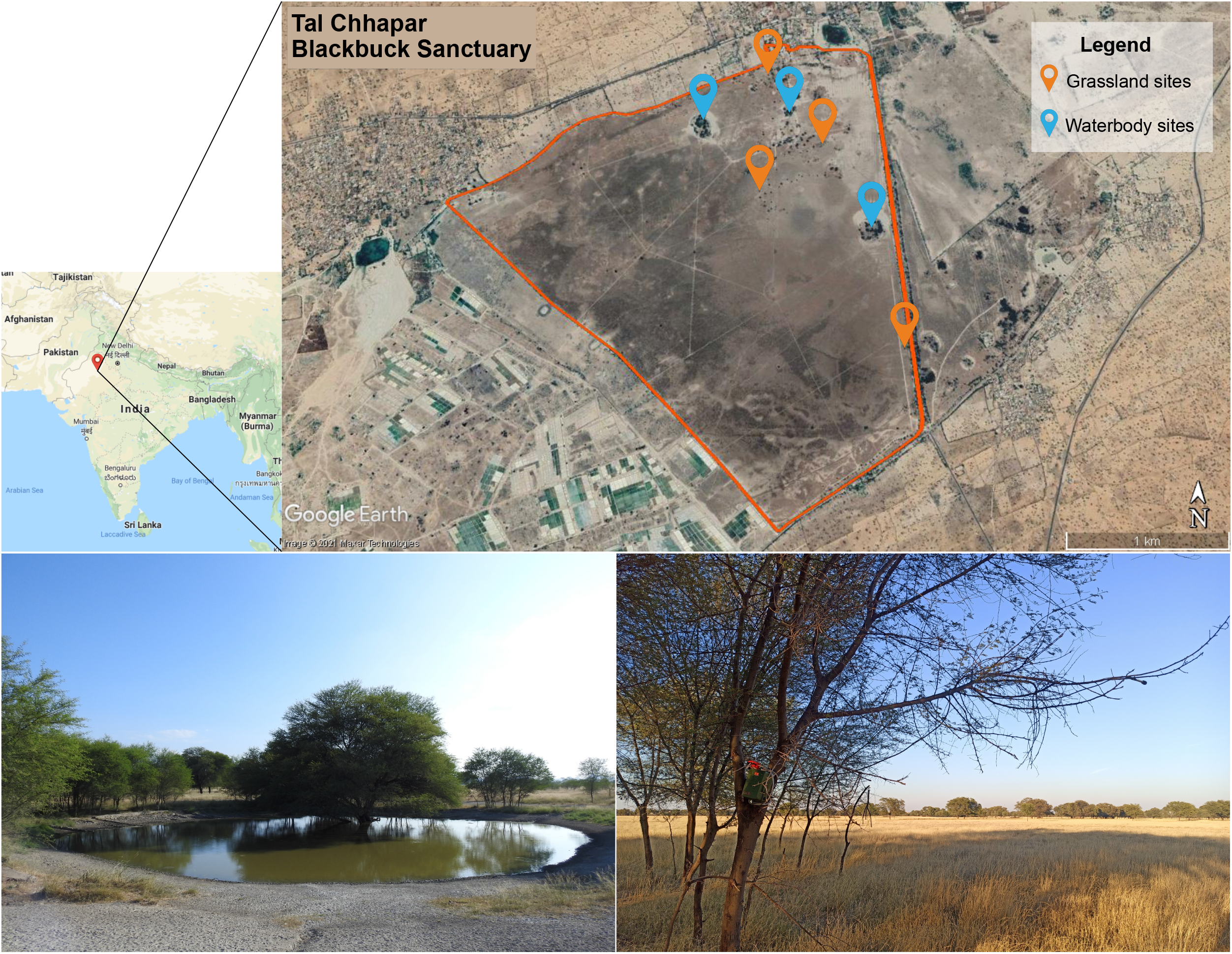
Map of the study area, Tal Chhapar Blackbuck Sanctuary in Churu district, Rajasthan, along with the locations of the sampling sites. The water bodies are marked in green and the grassland sites in red. Images on bottom left and bottom right are of a water body and a grassland recording site within the sanctuary.

### Passive acoustic sampling

Using acoustic recorders, we collected species-level information on bat activity and species composition for the bat assemblage at Tal Chhapar. Data were collected in November 2019, March 2020 and January 2021 during the winter dry season. This ecosystem experiences pronounced dry and wet seasons, and the portion of the dry season we sampled corresponded to low rainfall and changes in temperature (the winter), and is thus a good time to monitor changes in bat activity. Using SongMeter SM4BAT recorders (Wildlife Acoustics, Maynard, MA, USA), we recorded bat activity across 20 sampling nights (4 nights each at 3 water bodies and 2 nights each at 4 grassland sites) in November 2019. In March 2020 we collected 16 sampling nights of data (4 nights each at 3 water bodies and 1 night each at 4 grassland sites), because the COVID-19 lockdown terminated all fieldwork before we could reach 20 sampling nights. In January 2021 we collected another 20 sampling nights of data. Thus, our overall dataset contained 56 sampling nights of data across 7 sites over 3 seasons. Because the number of bat detections in January 2021 was too low for statistical analysis (Supplementary Figure 1), we performed most statistical comparisons on the data from November 2019 (early winter) and March 2020 (late winter), a total of 36 sampling nights. To record bat activity at water bodies, we deployed the recorders on trees near their periphery. At grassland sites, we deployed the recorders on trees at dispersed locations in the grassland.

During each sampling night (per recorder), we recorded 12 hours of acoustic data between 18:00 and 06:00 hrs (Muthersbaugh et al., 2019), totaling 672 sampling hours of recording time across all seasons, of which 432 sampling hours from November 2019 and March 2020 were used for further analysis. The audio settings of the recorders were set to a gain of 12 dB, sampling rate of 384 kHz, and triggered when they detected calls with a minimum duration of 1.5 milliseconds and a minimum trigger frequency of 10 kHz, with a trigger level of 12 dB and a trigger window of 1 second. We identified bat species from their echolocation calls based on their unique spectral structure and frequency parameters, using published information on bat calls from India and records of species known for the region (Chakravarty et al., 2021, 2020; Raman and Hughes, 2021; Shah and Srinivasulu, 2020; Srinivasulu et al., 2013; Wordley et al., 2014) as a reference, and verified this using field observations. For the latter, we combined observations of bats visiting waterholes before dark with recordings using a Wildlife Acoustics Echo Meter Touch 2 Pro mobile bat module plugged into an Android device. The module was set to a sampling rate of 384 kHz on live mode for active detection of bat calls.

### Temporal activity patterns

First, we subsampled our data by selecting six 5-minute samples (total 30 mins) spanning each hour of recording (0-5 minutes, 10-15 minutes, 20-25 minutes, and so on for each hour, using the time stamps on the recorded files). We used Raven Pro 1.5 (Cornell Laboratory of Ornithology, Ithaca, NY, USA) to visualize spectrograms from each subsample, using a Hanning window of size 512 samples with 50% overlap between windows. If at least two consecutive echolocation pulses of a sequence were present in a 5-minute sample, we marked the species’ presence by ‘1’, and conversely the absence with a ‘0’. Thus, we constructed a presence/absence (P/A) matrix for all species for each sampling night (72 samples per sampling night, spanning the entire 12-hour period) (Krishnan, 2019; Lahiri et al., 2021). We chose only search phase calls as a criterion for detection, because their acoustic parameters are relatively invariant within-species, and are thus more easily identifiable. Our metric (presence-absence in time windows) differed from the more commonly used metric of counting bat passes (Miller, 2001). However, by using presence-absence in given time windows, we were able to obtain conservative and directly comparable estimates of activity patterns for different species.

From the P/A matrix we calculated the frequency of detection for each species within each hour, which we defined as the fraction of 5-minute samples in which the species was detected, standardized to range between 0 to 1. Here, detection refers to the presence of a bat in a given sample. Thus, a species detected in all six 5-minute samples in any given hour, would have a detection frequency of 1 for that particular hour, whereas one detected in three samples would have a detection frequency of 0.5. We used these P/A matrices to determine the average detection frequencies for all 12 hours of the night in a given sampling season. This value served to compare both diel and seasonal bat activity.

We used circular statistic metrics to quantitatively describe diel activity patterns. Using Oriana 4.02 (Kovach Computing Services, Anglesey, UK), we conducted vector strength analyses to quantify circular dispersion in the temporal activity distributions of each bat species (Cremers and Klugkist, 2018). A relatively high mean vector length would indicate that the activity peaked at certain times of the night, while a lower mean vector length would be more consistent with relatively uniform activity levels across the night. We used the Rayleigh test of uniformity to assess the significance of mean vector lengths (Zar, 1999).

To compare detection frequencies across the seasons and habitat types, we conducted two-tailed Wilcoxon Rank-Sum tests in R (R Core Team, 2021) using the function ‘wilcox_test’, and calculated the effect sizes using the function ‘wilcox_effsize’ (Kassambara, 2021).

### Fine-scale temporal avoidance

Finally, we investigated fine-scale temporal avoidance in species with significant overlap in diel activity (specifically, *Pipistrellus* and *Scotophilus*, the most commonly detected species in our dataset). For this analysis, we used only data from water body sites, to test whether the two species partitioned arrival times at this resource. This analysis was conducted across the time span of 18:00 - 23:00 hours, as we determined that this was the duration in which both species were most active. Using Raven Pro, we visually examined each 1-minute file within a 5-minute sample (see above; each 5-minute sample surveyed consisted of 1-minute files representing each bat detection) for the presence of both species. When search phase calls of both species were detected simultaneously (i.e. the second species was detected while the first was still present), the event was considered an “overlap” in time (Supplementary Figure 2). On the other hand, the occurrence of only one of the two species within a one-minute sample, or the presence of both species without temporal overlap (i.e. the second species arrived after calls of the first were no longer detected) was considered a “non-overlap” event.

To examine whether coexisting bat species exhibited fine-scale temporal avoidance, we first calculated the ‘observed overlap’ values for each sampling night, or the total number of overlap events across a sample night. Our goal was to test whether the two species exhibited temporal avoidance on nights when both were active together. The probability of overlap expected by chance (‘expected overlap’) was calculated as *p_A_p_B_*, where *p_A_* and *p_B_* are the detection frequencies of *P. tenuis* and *S. heathii* respectively (Planqué and Slabbekoorn, 2008). This probability was multiplied by the total number of 1-minute segments analyzed, to determine the ‘expected overlap’ counts. For this analysis, we excluded three sampling nights where *S. heathii* was not detected, as both observed and expected overlap would be 0 in those cases. We compared observed and expected temporal overlap using a Wilcoxon signed-rank test in R, using the function ‘wilcox.test’. An observed temporal overlap that was significantly lower than expected by chance would indicate temporal avoidance between the two species.

## RESULTS

### Bat acoustic assemblage

We detected six species of bats in the bat assemblage of Tal Chhapar Sanctuary. The bat species found in the study area possessed unique call frequencies which enabled identification from acoustic data. We identified these call types as belonging to *Pipistrellus tenuis, Scotophilus heathii, Tadarida aegyptiaca, Rhinolophus lepidus, Rhinopoma hardwickii* and *Taphozous nudiventris*. (Figure 2A). In the case of *P. tenuis* and *S. heathii*, we verified species identity by observing bats and recording them using a mobile bat detector (see Methods). Call frequencies, which have been published as identification keys in previous studies on Indian bats, coupled with published information on bats known to occur in Rajasthan (Chakravarty et al., 2020; Raman and Hughes, 2021; Shah and Srinivasulu, 2020; Srinivasulu et al., 2013; Wordley et al., 2014), were additionally used to confirm identification. *S. heathii* was also identified by its yellow ventral pelage color, which is visible in the field (Srinivasulu et al., 2010). *Rhinopoma hardwickii* was detected only once in November 2019, whereas *Taphozous nudiventris* was detected once each in November 2019 and March 2020. Further analyses therefore focused on the other four bat species. Bat activity was very low in January 2021 compared to November and March (Supplementary Figure 1), and so we excluded this data from statistical analysis. The decrease in bat detections was, however, consistent with reduced activity during the peak of winter, followed by an increase in the warmer weather in March. Across site types, we observed a significantly higher number of bat detections near water bodies compared to grassland sites (W=15.5, p=1.7e-5, r=0.719) (Figure 2B). This is consistent with water body monitoring being a more effective tool to detect bats in these habitats, particularly as they descend to drink. However, given the relatively small size of this sanctuary, our analysis of activity patterns used pooled data across sites, as bats were likely to be transiting across the grassland sites to visit the water bodies.

**Figure 2:**
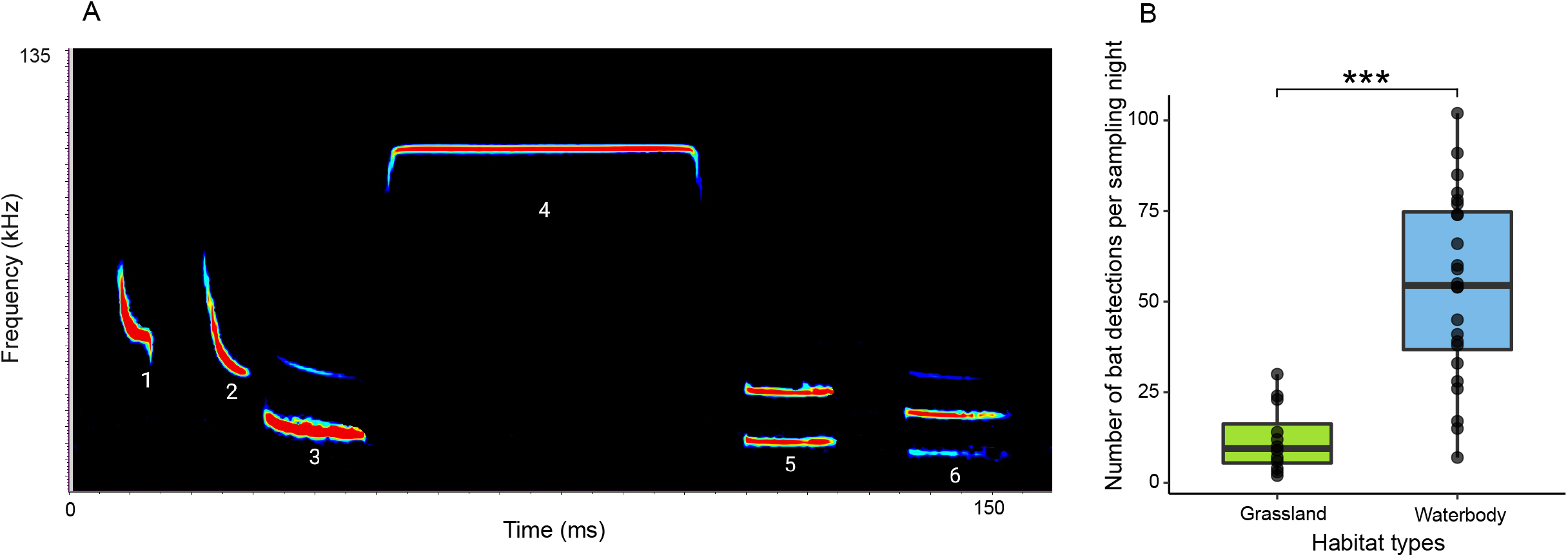
**A**: Composite spectrogram of bat echolocation calls from the study area. The calls depicted are : (1) *Pipistrellus tenuis*, (2) *Scotophilus heathii*, (3) *Tadarida aegyptiaca*, (4) *Rhinolophus lepidus*, (5) *Rhinopoma hardwickii*, (6) *Taphozous nudiventris*.**B**: Box plot comparing the mean bat detections between grassland sites and water bodies.

### Activity patterns

The nocturnal activity patterns of the four species of bats (representing pooled data across all sites, and thus a combined activity pattern measure for the entire sanctuary) are presented in Figure 3. *Pipistrellus tenuis* was active throughout the night (mean vector length = 0.712; Rayleigh test: Z = 238.1, p<0.001, n = 470), with a peak in activity between 20:00-21:00 in both November and March. On the other hand, *Scotophilus heathii*’s activity patterns exhibited a distinct peak during early evenings (mean vector length: 0.854, Rayleigh test, Z = 111.6, p<0.001, n = 153), between 18:00-19:00 in November and 19:00-20:00 in March, and activity levels decreased considerably following this peak. The other two bat species, *Tadarida aegyptiaca* and *Rhinolophus lepidus*, were detected relatively rarely compared to the previous two species but were active throughout the night (*T. aegyptiaca*: mean vector length = 0.827, Rayleigh test, Z = 20.6, p<0.001, n = 30); *R. lepidus*: mean vector length = 0.716, Rayleigh test, Z = 6.2, p = 0.001, n = 12).

**Figure 3:**
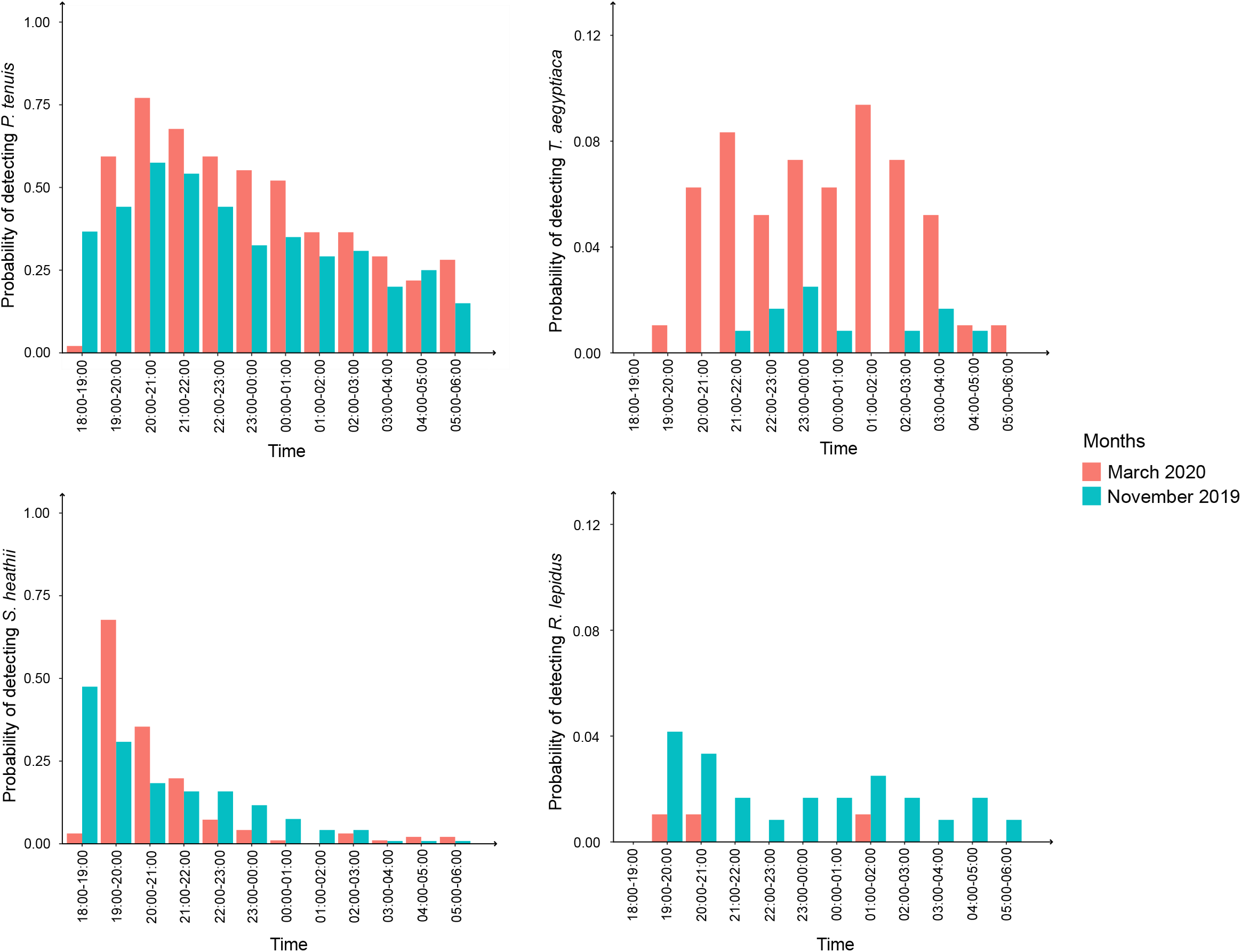
Overnight activity patterns of *P. tenuis* (top left), *S. heathii* (bottom left), *T. aegyptiaca* (top right) and *R. lepidus* (bottom right) in November 2019 (blue) and March 2020 (red). Here, the probability of detecting a species (in y-axis) ranges from 0 to 1 for all species.

### Seasonality in bat activity

*P. tenuis* and *S. heathii* were detected at similar levels in both November (n = 20 nights) and March (n = 16 nights) (*P. tenuis*: W = 133.5, p =0.408, r = 0.141; *S. heathii*: W = 152.5, p = 0.823, r = 0.040). On the other hand, *Tadarida aegyptiaca* and *Rhinolophus lepidus* exhibited seasonal changes in activity between November (n = 20 nights) and March (n = 16 nights) (*T. aegyptiaca*: W = 25.5, p = 1.09e-5, r = 0.736; *R. lepidus*: W = 221.5, p = 0.020, r = 0.390). *T. aegyptiaca* was detected more frequently in March, whereas *R. lepidus* was detected more frequently in November and very rarely in March. Overall, the total number of detections for all species combined did not differ significantly between November and March (W = 126, p = 0.286, r = 0.180). Coupled with the lower detections in January, this is consistent with seasonal turnover in the bat assemblage, with differences in species abundance after the peak of winter.

### Fine-scale temporal partitioning of bats at scarce water resources

Our data on nocturnal and seasonal activity patterns suggested that bats in these semiarid grasslands exhibit species-specific activity patterns. The two most common species, *P. tenuis* and *S. heathii*, however, were active through the seasons covered in our sampling period, and both were very active in the early evening, following which *S. heathii* declined through the night (Figure 3). We therefore examined if the two species exhibited temporal overlap at finer time scales during this early evening period when visiting water bodies. The observed overlap values between these two species were significantly lower than the overlap values expected by chance (Figure 4) (W = 231, p = 9.54e-7, r = 0.88). This is consistent with the two species exhibiting temporal avoidance when their diel activity patterns overlap.

**Figure 4:**
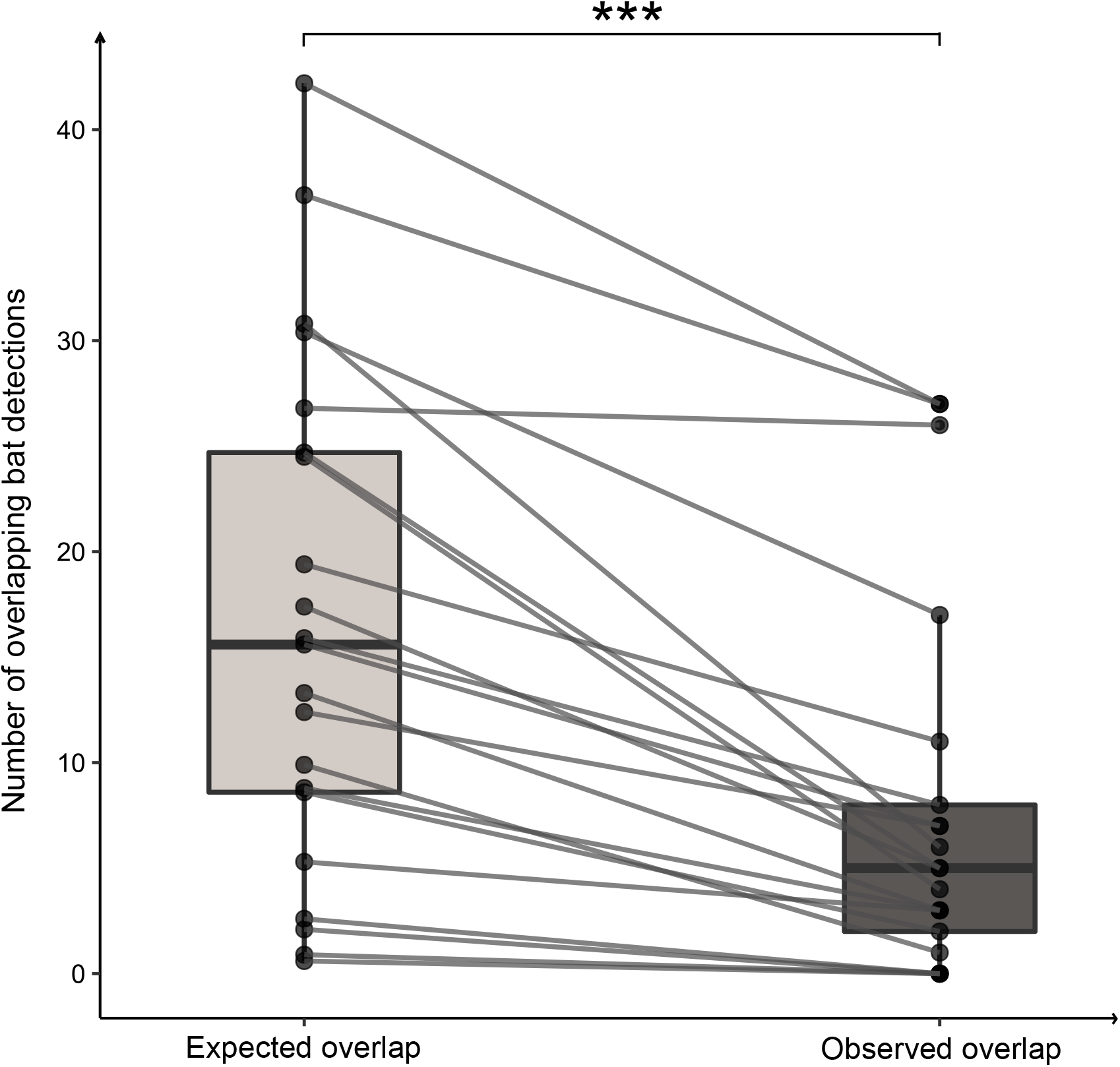
Observed temporal overlap in detections of *P. tenuis* and *S. heathii* is much lower than expected by chance.

## DISCUSSION

In summary, we find that bats in semiarid grasslands were much more easily detected at sources of water, and that some bat species exhibit constant activity patterns and detection rates through the winter dry season whereas others exhibit variation as the season progresses. Finally, we find that the two most frequently detected species coexist by temporally partitioning their use of a scarce and potentially limiting resource (water), by alternating their arrival even at fine time scales. Thus, the two species minimize overlap at this resource even though their peak activity periods significantly overlap. Our data thus identifies temporal patterns in the structure of the bat assemblage occupying this habitat, and we discuss these patterns and their implications below.

### Temporal partitioning at scarce resources

In diverse animal taxa, competition for scarce or patchily distributed resources may drive behavioral strategies that promote coexistence, while minimizing competition. For example, divergent traits or spatial segregation of foraging and vocal activity help minimize competitive overlap (Begon and Townsend, 2020; Chitnis et al., 2020; Jain and Balakrishnan, 2012; Krishnan and Tamma, 2016; MacArthur, 1958). In addition, many organisms are temporally partitioned, with peak activity at different times of the 24-hour diel cycle (Hofmann et al., 2016; Kemna et al., 2020; Luther, 2008; Ullas Karanth et al., 2017). Birds partition singing activity to minimize temporal overlap, by either singing at different times of the day or by timing their songs to avoid overlap at fine scales (Chhaya et al., 2021; Cody and Brown, 1969; Planqué and Slabbekoorn, 2008; Popp et al., 1985). Here, we uncover evidence that the two most frequently detected bat species in the Tal Chhapar landscape, *Pipistrellus tenuis* and *Scotophilus heathii*, temporally partition so as not to overlap at scarce water resources. Although their peak diel activity levels overlap extensively with each other, the two species alternate their arrivals at water bodies such that observed overlap is much less than expected by chance. Even where the two arrive within one minute of each other, one species typically does not arrive until the other has departed. Both species echolocate at different frequencies (Figure 2), so acoustic interference is unlikely to drive this avoidance behavior.

Water is a scarce resource in arid landscapes, and multiple studies have demonstrated its importance in maintaining bat assemblages within such habitats (Loumassine et al., 2020; Racey et al., 1998; Williams et al., 2006). Partitioning of arrival times or arrival paths at water holes is thought to minimize overcrowding or in-flight collisions, and may be a key driver of species coexistence in relatively extreme environments (Adams and Simmons, 2002). *P. tenuis* and *S. heathii* are present at roughly similar detectability levels throughout the dry season, and the observed temporal partitioning thus likely permits the smaller species to coexist with the larger one even in conditions of water scarcity. We note here that *P. tenuis*, which possesses higher vocal frequencies, was detected more often than *S. heathii*. This, together with the small size of the water bodies at which we made these measurements, suggests that differences in detection distance did not influence the patterns we observed. As climate change is predicted to result in increasing aridity of these environments, coexistence mechanisms such as these may support the resilience of bats to water scarcity. Long-term monitoring studies will help elucidate these patterns, as we discuss below.

### Seasonality of bats in arid environments

Diverse animal taxa survive in extreme environments by undergoing seasonal changes to their physiology or movement patterns. Any of these seasonal behavioral changes may result in a change in community structure, with the absence of certain species from a landscape. Bats, as one of the most speciose orders of mammals, exhibit complex responses to seasonal weather patterns. These responses include migration to more favorable climates, metabolic adjustments to remain active, and hibernation (Fleming et al., 2003; Geiser, 2013). Animal seasonality is thought to be a response to changes in food availability, and is thus driven by energetic considerations, with temperature and photoperiod as likely initiating cues (Meyer et al., 2016; Williams et al., 2014). Although seasonality of bats in temperate regions is well-documented (Heim et al., 2016; Loumassine et al., 2020), we still lack comprehensive data on the seasonal dynamics of bat assemblages in tropical arid landscapes, particularly in South Asia where bats as a whole are very poorly studied. The extreme temperatures and water scarcity in areas such as the Thar desert renders them good places to study and monitor these seasonal movements.

Indeed, we find evidence of pronounced seasonality in bats inhabiting semiarid grasslands. *Tadarida aegyptiaca* was detected much more frequently in March, whereas *Rhinolophus lepidus* was detected much more frequently in November. This suggests either some partial migration to different sites, or a change in overall activity levels, potentially driven by changes in temperature or prey availability. *Rhinopoma hardwickii* is a species in our dataset which is known to hibernate; we recorded this species only once during our sampling in the sanctuary, but found a colony of hibernating bats of this species in a nearby village. It is possible that this species does not utilize grassland habitats, or visits larger water bodies than those present in the sanctuary, and further studies are needed to examine this. Finally, we found that detections of all bat species were very low during January, when temperatures are near their lowest point in the year. This is consistent with our interpretation of seasonal turnover in the bat assemblage, in that bat activity drops, and then recovers by March, but with different abundances of the various bat species. Additionally, this observation suggests that bat monitoring should be concentrated during the early and late parts of the winter season, when bat activity levels are higher.

### Monitoring bat activity patterns in semiarid grasslands

Arid regions of the world are commonly neglected in conversations about the effects of climate change, land-use change and conservation in general. Because policy decisions in many parts of the world are made considering these habitats as “wastelands”, their biodiversity has been neglected in monitoring efforts (Joshi et al., 2018). The sensitivity of bats to environmental changes renders them excellent indicators of ecosystem health (Jones et al., 2009). Studies that seek to monitor seasonality, activity patterns and abundance are of great importance in establishing a baseline for long-term monitoring. Water bodies are extremely important in monitoring bat assemblages in arid regions, as they are crucial to support bat diversity in extreme environments (Racey et al., 1998). We find that bat detections at water bodies are considerably higher than those at sites in the grasslands, concordant with other studies that uncover the same pattern, particularly during the dry season when water and other resources are scarce (Barros et al., 2014; Lisón and Calvo, 2011; Shapiro et al., 2020; Vaughan et al., 1997; Williams et al., 2006). Bats descending to drink at water bodies are more likely to be detected by bat recorders, and bats detected in the grasslands may have been transiting to and from these water bodies. For this reason, we present a general activity pattern rather than examining activity differences across sites. Thus, our pooled activity pattern measurements across all sites represent a generalized diel pattern for the entire sanctuary. A paradigm in long-term monitoring of bats in this landscape should therefore focus on monitoring bats at the few water sources that exist in the dry season, to effectively census the community and activity patterns.

The long-term effects of climate change are predicted to result in increasing aridification of deserts and semiarid landscapes (Mirzabaev et al., 2019). Thus, the already scarce (and potentially limiting) water resources are likely to become more unpredictable, with consequences for the entire ecosystem. Because bats are concentrated at water sources and are so sensitive to environmental changes, increasing aridity may influence both seasonal movement patterns of bats, as well as fine-scale temporal partitioning. By quantifying and demonstrating that both these patterns exist in the bat assemblages occupying these landscapes, we offer both a demonstration of the complex temporal dynamics of bat assemblages in semiarid grasslands, as well as their potential to detect the effects of climate change. Species such as *P. tenuis* and *S. heathii*, which coexist at limited sources of water, may also be able to inform us about the resilience of bats to water scarcity. These, and other trends, are best studied by a comprehensive, long-term monitoring program, for which we hope our study will serve as a launching point.

## Acknowledgments

We are grateful to the Rajasthan Forest Department, specifically the Chief Wildlife Warden (Shri Arindam Tomar) for the research permits (Permit letter number: F19()permission/cwlw/2017/1598) to conduct research at Tal Chhapar Wildlife Sanctuary, as well as the DCF of Churu District, ACF Tal Chhapar (Mr. Pradeep Chaudhary), forest department officers, Mr. Umesh Bagvatiya and Mr. Yogendra Singh Rathore for their invaluable help. We also thank Mr. Rishpal and other staff at the Tal Chhapar Sanctuary for their hospitality and assistance during sampling. Finally, we thank Arpit Omprakash, Sutirtha Lahiri and Varun Kher for help during sampling, M. Abhinava Jagan for comments on the manuscript and Rohit Chakravarty for input on sampling design and species identification.

---

**Supplementary Figure 1:** Differences in bat detections across seasons (November 2019, March 2020 and January 2021). Each data point represents a single sampling night.

**Supplementary Figure 2:** Example spectrogram of an “overlap” event between *P. tenuis* and *S. heathii* at a water body.

**Supplementary Data:** Supplementary Data file.

