## Supplementary figures and images for "Seasonality and interspecific temporal partitioning in a semiarid grassland bat assemblage"

### Supplementary Figure 1

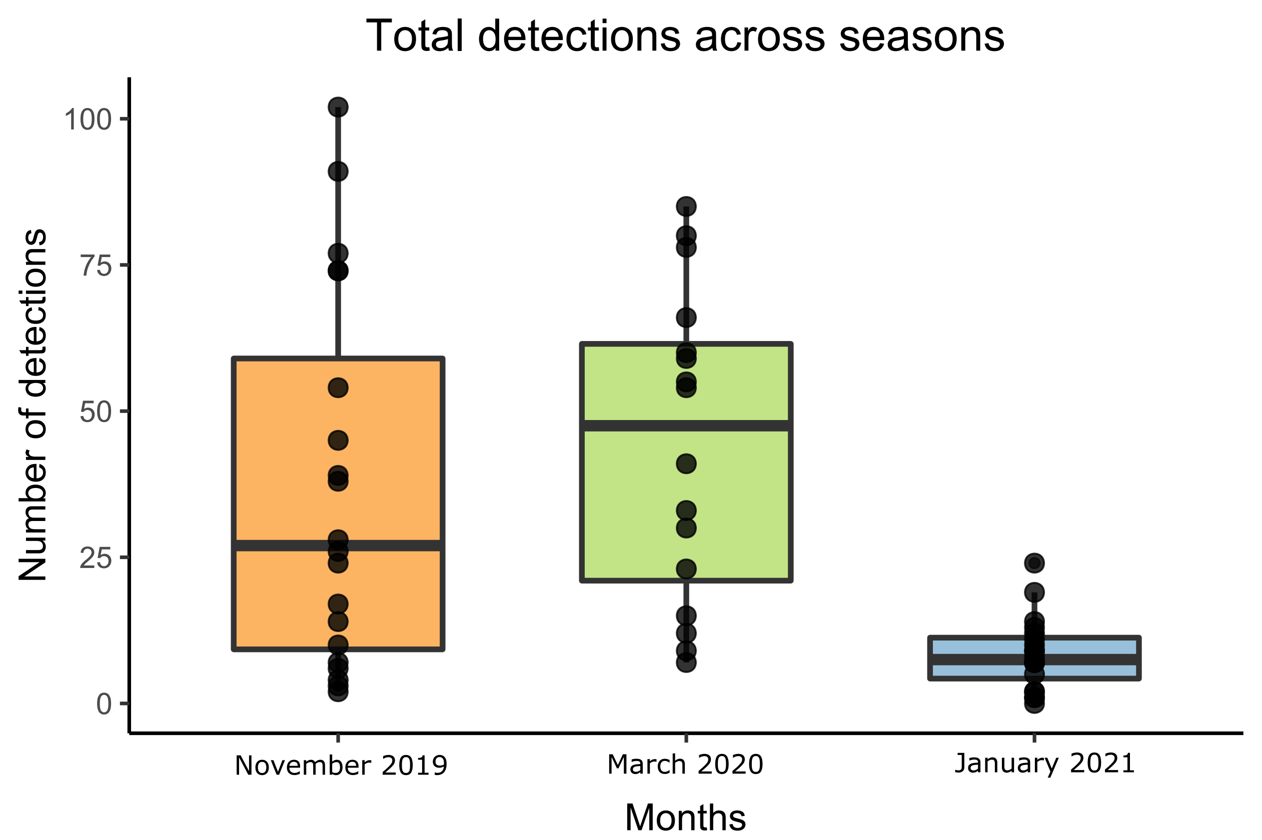

### Supplementary Figure 2

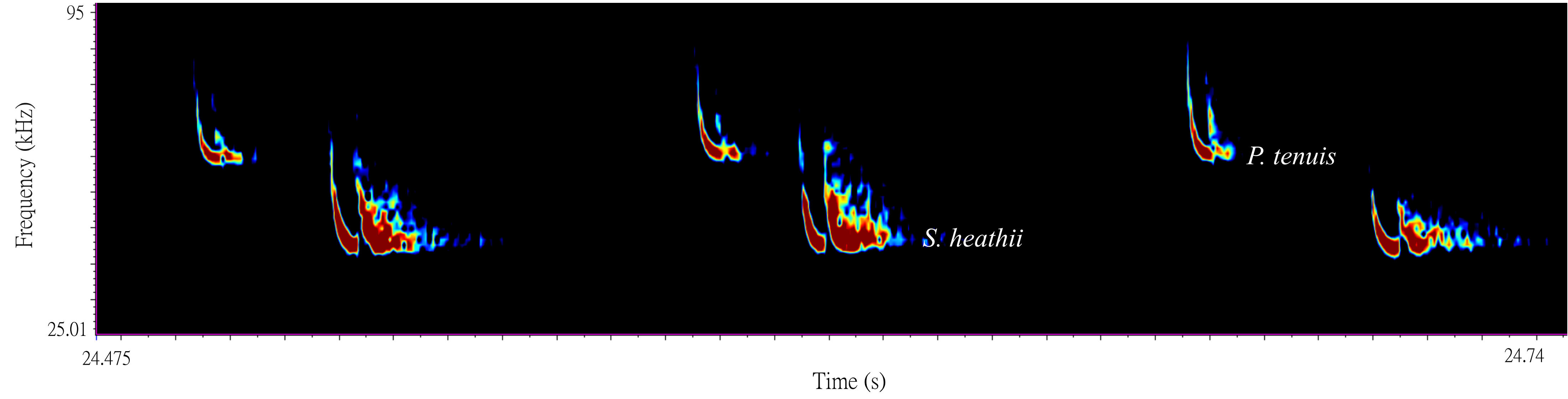
